# Sex specific regulation of the cortical transcriptome in response to sleep deprivation

**DOI:** 10.1101/2023.12.05.570075

**Authors:** Tianyi Shi, Ishani Shah, Quang Dang, Lewis Taylor, Aarti Jagannath

## Abstract

Multiple studies have documented sex differences in sleep behaviour, however the molecular determinants of such differences remain unknown. Furthermore, most studies addressing molecular mechanisms have been performed only in males, leaving the current state of knowledge biased towards the male sex. To address this, we studied the differences in the transcriptome of the cerebral cortex of male and female C57Bl/6J mice after six hours of sleep deprivation. We found that several genes, including the neurotrophin growth factor *Bdnf*, immediate early genes *Fosb* and *Fosl2*, and the adenylate cyclase *Adcy7* are differentially upregulated in males compared to females. We identified the androgen-receptor activating transcription factor EZH2 as the upstream regulatory element specifying sex differences in the sleep deprivation transcriptome. We propose that the pathways downstream of these transcripts, which impact on cellular re-organisation, synaptic signalling and learning may underpin the differential response to sleep deprivation in the two sexes.

## INTRODUCTION

Sleep is fundamental to many biological processes including metabolism, immunity, and memory formation. The mechanisms of sleep control are not fully understood, but it has been posited that sleep is controlled by a homeostatic process (Process S) and a circadian process (Process C), that interact to regulate sleep/wake cycles (Borbély, 1982; Borbély et al., 2016). Process S, represented as sleep drive, depends on the intensity and duration of the preceding period of wakefulness. Whereas Process C is controlled by the master circadian pacemaker/clock located in the suprachiasmatic nucleus (SCN), for which the key zeitgeber is light-information received from intrinsically photosensitive retinal ganglion cells (ipRGCs) (Eban-Rothschild et al., 2018).

Circadian and sleep disruption can impact health, including cognitive function in mammals (Simon et al., 2022). The transcriptomic changes associated with the homeostatic response to sleep deprivation have been examined in rodents using microarrays and RNA-seq with results showing that immediate early genes (IEGs) Homer Scaffolding Protein 1 (*Homer1*), Activity Regulated Cytoskeleton Associated Protein (*Arc*), and brain-derived neurotrophic factor (*Bdnf*) are consistently upregulated (Maret et al., 2007). Sleep deprivation has also been shown to regulate circadian clock genes with upregulation of *Period ½ (Per1/2)* and downregulation of D-Box Binding PAR BZIP Transcription Factor (*Dbp*) (Cirelli et al., 2004; Gerstner et al., 2016; Mackiewicz et al., 2007; Terao et al., 2006; Thompson et al., 2010; Wisor et al., 2008). However, one caveat regarding these studies is that they only used male rodents. Under-representation of female rodents in neuroscience research is common due to the notion that inclusion of female rodents would lead to greater variability in comparison to male rodents. However, one meta-analysis looking at 9,932 traits has disputed this assumption, highlighting how there is no substantial difference between female and male variability (Prendergast et al., 2014). This misconception has therefore limited our understanding of sex differences, across the field of neurobiology, and more specifically within sleep and circadian research.

In sleep and circadian research, there is growing evidence for sex differences in sleep and circadian physiology in both humans (Carrier et al., 2017; Manuel Spitschan et al., 2022; Mong & Cusmano, 2016; Santhi et al., 2016) and murine models (Dib et al., 2021a). In humans, these sex differences in sleep can be summarised as women reporting poorer sleep quality with more frequent insomnia symptoms than men (R. H. Y. Li et al., 2002; Mong et al., 2011). However, quantitative studies have shown that women exhibit an overall longer sleep time than men, with EEG results showing women have higher EEG power in most frequencies during the two main sleep states (non-rapid eye movement (NREM) and rapid eye movement (REM)) suggesting higher quality NREM sleep and lower quality REM sleep (Carrier et al., 2001; Dijk,’ et al., 1989; Long et al., 2021). These differences are dependent on other factors including age, and phase of estrous cycle, and species. In mice, studies indicate that males exhibit reduced wakefulness, but when in proestrus, females show increased wakefulness compared to males during the light and early dark period (Swift et al., 2020). Rodent sleep studies investigating sex differences have demonstrated that gonadectomy (GDX) eliminates sex differences in wakefulness by significantly decreasing the disparity in females, thus suggesting ovarian hormones contribute to time spend sleeping in rodents with female-specific brain organisation (Cusmano et al., 2014; Paul et al., 2006). EEG studies in mice provide conflicting results as to whether males spend more or less time in NREM sleep than females (Choi et al., 2021; Nichols et al., 2020; Paul et al., 2006), but the majority report higher NREMS sleep in males (Dib et al., 2021a). Further studies focusing on the contribution of sex chromosomes demonstrate that males with either XX or XY chromosomes spend more time in NREM than females with XX or XY chromosomes, suggesting gonadal hormones regulate NREMS (Nichols et al., 2020; Ehlen et al., 2013).

Despite this evidence, our mechanistic understanding of these differences is poor. Current research suggests the involvement of both hormone-dependent and hormone independent elements (Paul et al., 2009). Previous rodent studies using acute sleep deprivation (6-8 hours) found that females exhibit either a stronger (Paul et al., 2006) or weaker (Choi et al., 2021; Nichols et al., 2020) NREM rebound in the recovery sleep, under different experimental settings. Again, these differences were reduced by GDX (Paul et al., 2006), providing more support for ovarian hormone contribution to sex differences in homeostatic sleep regulation. To explore the transcriptomic basis accountable for these differences, we performed RNA-seq on the cerebral cortex of acutely sleep-deprived mice of both sexes.

## METHODS

### Resource Availability

#### Lead contact

Further information and requests for resources and reagents should be directed to and will be fulfilled by the lead contact, Aarti Jagannath *Materials availability:* This study did not generate new unique reagents.

#### Data and code availability

All RNA-Seq data have been deposited on NCBI SRA and will be publicly available as of the date of publication. No original code was used in this study. Any additional information required to reanalyse the data reported in this paper is available from the lead contact upon request.

### Experimental Model and Subject Details

#### Animals

All studies were conducted using male and female C57BL/6 J mice over 8 weeks of age and, unless otherwise indicated, animals were group housed with *ad libitum* access to food and water under a 12:12 hour light/dark cycle (100 lux from white LED lamps). All animal procedures were conducted in accordance with the UK Home Office regulations (Guidance on the Operation of Animals (Scientific Procedures Act) 1986) and the University of Oxford’s Policy on the Use of Animals in Scientific research, following the principles of the 3Rs.

### Apparatus and Experimental Procedures

For the Sleep Deprivation (SD) experiments, animals were kept awake for 6 hours between ZT0 (Zeitgeber time 0 or lights on within the 12h/12h light/dark cycle) and ZT6 (Zeitgeber time 6 or 6h into the light period within the 12h/12h light/dark cycle) by providing novel objects to elicit exploratory behaviour, as previously described (Huber et al., 2000). The animals were then sacrificed (4 males and 4 females per condition), and somatosensory cortical tissue punches collected. Control animals were allowed to sleep ad libitum between ZT0 and ZT6. Variation in sleep/wake and circadian behaviour through the oestrous cycle (Dib et al., 2021b) (Joye & Evans, 2022) is often a reason to exclude female mice during experiments and oestrous cycle monitoring can be cumbersome, which can lead many experimentalists to avoid using females. We did not monitor oestrous cycle, in order to capture variation as would be expected if oestrous stage is not controlled.

### Method Details

#### RNA extraction and RNA sequencing library preparation

Total RNA from cortical punches was extracted using TRIzol and the RNeasy Mini Kit (Qiagen). Cortical tissue was mechanically disrupted in 700 µl of TRIzol and 140 µl of chloroform was added and the sample thoroughly mixed. Following a 3 min incubation at RT, the sample was then centrifuged for 15 min at 15,000 xg, 4°C. The clear top layer was then carefully collected, mixed with an equal volume of 70% ethanol and RNA extracted using the RNeasy Mini Kit, with on-column DNase digestion, following the manufacturer’s instructions. RNA was eluted in water and RNA concentration and quality were measured using a TapeStation system (Agilent) with the High Sensitivity RNA ScreenTape assay. mRNA purification and cDNA synthesis for the sequencing library were performed according to the Illumina Stranded mRNA Prep protocol (20040534) using the following index kit: IDT for Illumina RNA UD Indexes Set A, Ligation (20040553). Quality and concentration of the final libraries were checked with the KAPA Library Quantification Kit (Roche Diagnostics) in a StepOnePlus thermal cycler (Applied Biosystems) according to manufacturer’s instructions. All cDNA libraries were sequenced using a paired-end strategy (read length 150 bp) on an Illumina NovaSeq platform.

#### Processing of RNA sequencing data

Raw RNA-Seq data processing (quality control, trimming, mapping to the genome, and read counting) was performed using tools embedded in Galaxy (v21.05) (Jalili et al., 2020). The fastqsanger files containing the raw sequencing data were uploaded to the public Galaxy server at usegalaxy.org. FastQC (v0.11.8) (https://www.bioinformatics.babraham.ac.uk/projects/fastqc/) was used for quality control of sequencing data. For quality and adapter trimming, Trim Galore! (v0.6.3) (https://www.bioinformatics.babraham.ac.uk/projects/trim_galore/) was employed to remove low-quality bases, short reads, and Illumina adapters. High quality reads were then mapped to the Mus musculus (mm10) reference genome using HISAT2 (v2.1.0) (D. Kim et al., 2019), specifying the strand information as reverse. featureCounts (v2.0.1) (Liao et al., 2014) was run to quantify the number of reads mapped to each gene. The featureCounts built-in mm10 gene annotation file was selected and under paired-end reads options, the option to count fragments instead of reads was enabled. The generated counts files were converted to CSV and downloaded for downstream differential gene expression analysis in R. MultiQC (v1.9 https://multiqc.info) (Ewels et al., 2016) was used to aggregate FastQC, HISAT2, and featureCounts results.

#### Differential gene expression analysis

To identify differentially expressed genes in males versus females under baseline conditions and after sleep deprivation, the DESeq2 package (v1.32.0) (Love et al., 2014) was used in R (v4.2.0, https://www.r-project.org). DESeq2 corrects for multiple testing using the Benjamini-Hochberg (BH) method, and only genes with a BH adjusted p value < 0.05 were considered statistically significant. To map differences in the response to sleep deprivation in the two sexes, we studied those genes that were differentially expressed in only one sex AND displayed a difference of greater that 0.5 log2-fold change after sleep deprivation. Heatmaps were drawn using the pheatmap function from the pheatmap package (v1.0.12). Volcano plots were generated using the ggplot2 package (v.3.3.5). STRING (database of known and predicted protein-protein interactions) interaction network mapping and functional annotation was conducted using string-db.org.

## RESULTS

A comparison of the sleep deprivation regulated transcriptome (Supplementary table 1) of the cortex in males versus female mice showed that broadly, the top 100 differential transcripts were similarly regulated in both sexes. For example, *Homer1, Dio2 (*Iodothyronine deiodinase 2*), Dusp4 (*Dual Specificity Phosphatase 4*), Cdkn1a (*Cyclin Dependent Kinase Inhibitor 1a*), Dbp* and *Cirbp (*Cold Inducible RNA Binding Protein), all shown previously to be regulated by sleep deprivation, changed consistently in both sexes (Figure 1A). 11 transcripts were identified as significantly different between males and females in a two-way comparison at baseline (Male Sham vs Female Sham) and 12 after sleep deprivation (Figure 1B). Many of these genes are on sex chromosomes (e.g. *Ddx3y, Eif2s3y* and *Eif2s3x*), but several were not – including the *Acer2* (alkaline ceramiase 2 which catalyses the hydrolysis of ceramides into sphingoid bases), *Lcat* (extracellular cholesterol esterifying enzyme lecithin-cholesterol acyltransferase), and *Scn7a* (voltage gated sodium channel protein type 7 subunit alpha), which lie on chromosomes 4, 8 and 2 respectively. These results demonstrate sex-specific gene regulation that is not related to the sex chromosomes.

**Figure 1:**
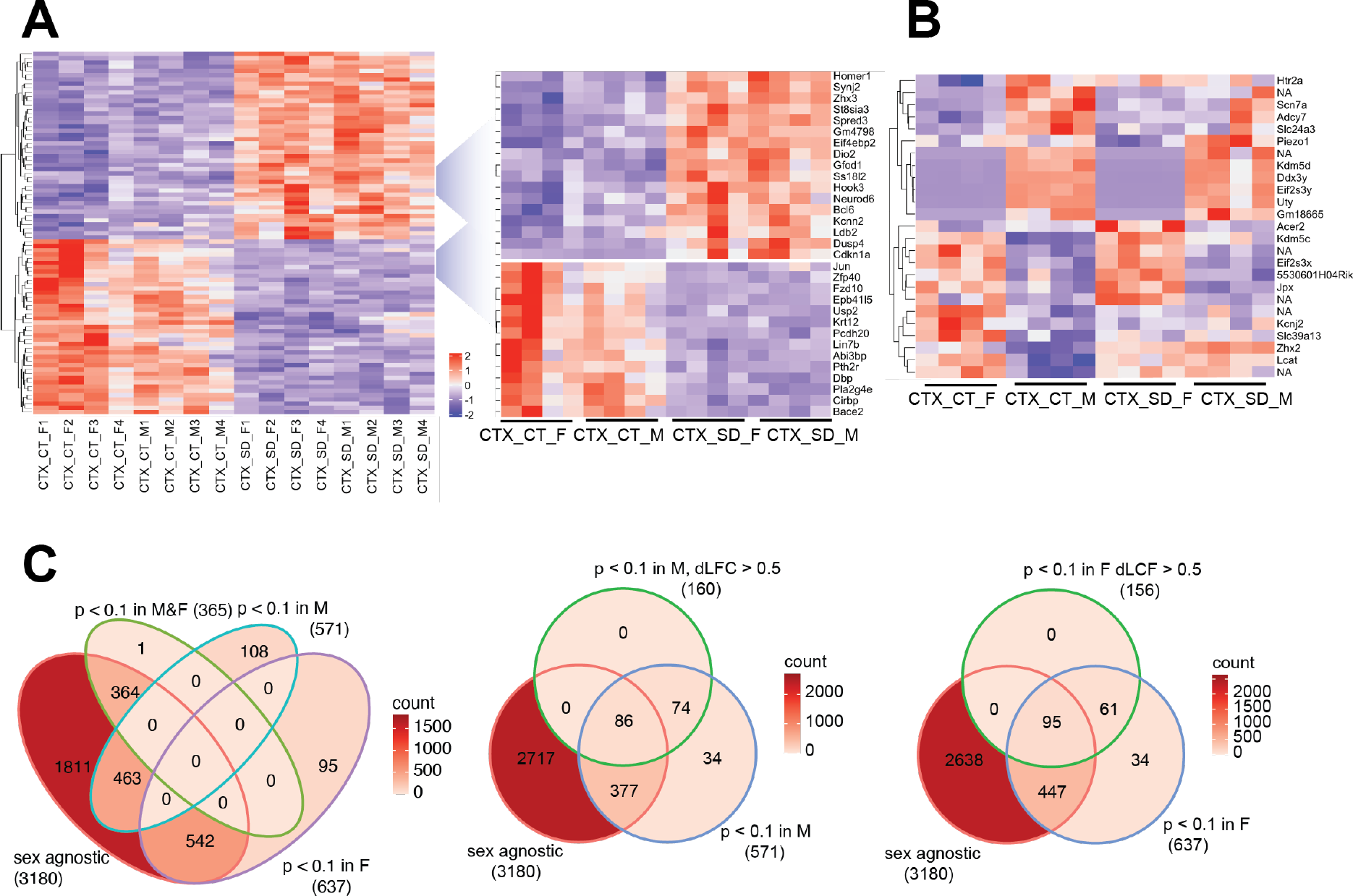
The transcriptional response in the cortex in response to sleep deprivation: (A) The top 100 differentially expressed transcripts in the cortex (CTX) under control conditions (CT) or 6h sleep deprivation (SD) in 4 males (M1-4) and 4 females (F1-4). Key upregulated and downregulated clusters highlighted on the right. (B) Transcripts that are significantly different in a direct comparison across the two sexes, either after sleep deprivation (SD) or under control (CT) conditions. (C) Venn diagrams showing the transcript numbers that are significantly different after sleep deprivation in the different conditions shown (sex agnostic, p adj < 0.1 in males only (M), females only (F), males and females (M&F). The central and right panel show those transcripts that are p adj < 0.1 in males only (M) or females only (F), but also differ between males and females by a log fold change of > 0.5 (dLFC>0.5). NA is non-coding transcript with no associated genes.

We next assessed the relative response to sleep deprivation (SD vs sham), between males and females. In total, 3180 transcripts were seen to change in response to sleep deprivation (p adj. <0.1) when sex was not included as a factor in the analysis. Of these, 364 were significant in both sexes, with approximately 600 genes significantly different in only one sex and not the other (Figure 1C). However, it is important to consider that the power of the experiment is reduced when analysing the two sexes separately (n=4 for each sex vs n=8 for both sexes). It is possible that the direction and general magnitude of change in both sexes is similar, but passes the statistical significance threshold for only one sex. To account for this, we identified those transcripts which were both significantly differential in only one sex and also different between the sexes by greater than 0.5 log2 fold change. In this manner, 316 transcripts were identified as changing in a sex-specific manner: 160 in males, and 156 in females (Figure 1C, Supplementary table 2). Several of these transcripts are illustrated in Figure 2A.

**Figure 2:**
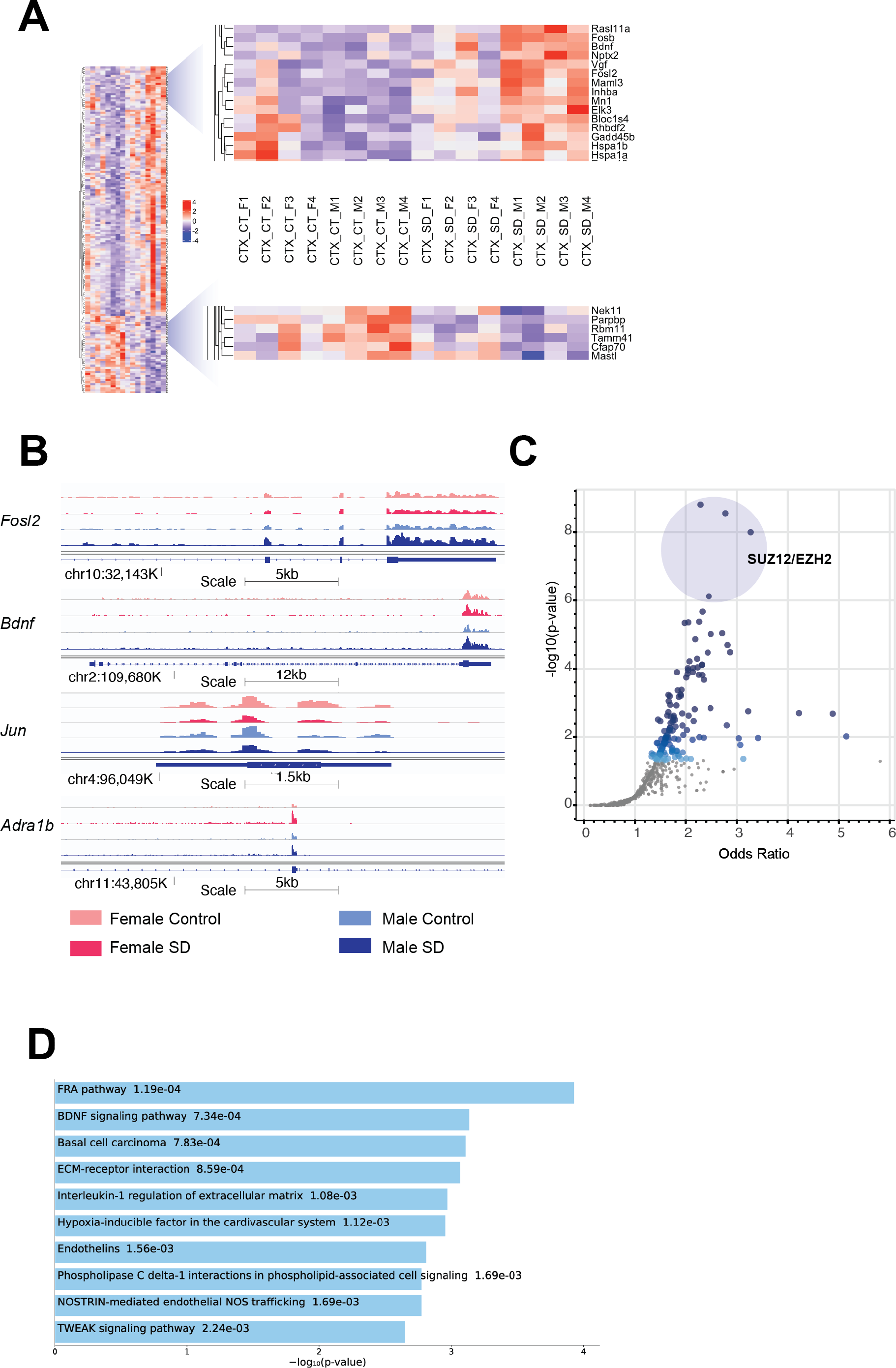
Sex-specific patterns of transcription in the cortex in response to sleep deprivation: (A) The transcripts with sex-specific regulation in response to sleep deprivation, i.e. p adj < 0.1 in males only (M) OR females only (F), and also transcripts that differ between males and females by a log fold change of > 0.5 (dLFC>0.5). Legend for samples as in Figure 1. (B) RNA-Seq tracks for genes of interest as labelled, genomic position and scale indicated. (C) Transcription Factor prediction for the transcripts in (A) using CHEA Transcription Factor Targets dataset on Enrichr. Binding sites for EZH2 and SUZ12 are significantly enriched, p-value indicated on Y-axis and X-axis the odds ratio (z-score assessing the rank of a term from a Fisher’s exact test against the expected rank of the term in the gene-set library. (D) Functional annotation of the transcripts in (A) from the Enrichr platform, Bioplanet 2019 results reported, the number alongside the term reports p-value.

Comparison between males and females shows that *Bdnf* (Brain Derived Neurotrophic Factor; key to plasticity, learning, and memory) and its targets (immediate early genes *Egr3, Fosb* and *Fosl2)* are upregulated more in males following sleep deprivation, as confirmed using pathway analysis (p = 9.3e^-5^) (Figure 2A, 2B). Increased transcription around exon 4 (Bdnf variant 4) is observed in both sexes after SD, but transcription from the coding exons in increased more strongly in males (Figure S2C). The same pattern of higher upregulation in males is seen with heat shock proteins (*Hsp1a* and *Hsp1b)* (Figure 2A). In contrast, *Jun*, and the adrenergic receptor *Adra1b* show significantly stronger downregulation and upregulation respectively in females (Figure 2B). Transcription factor *Jun* is one of the primary effectors of *HIF1a* (hypoxia inducible factor 1a), for which interestingly pathway analysis shows that other HIF signalling transcripts such as Nostrin (upregulated) and endothelin1/3 (downregulated) are differentially expressed in females following sleep deprivation, but not males. Other genes of interest include the RNA binding protein *Rbm11* which shows stronger downregulation in males (Figure 2A). Clock genes showed broadly similar responses to sleep deprivation as described previously, and in both sexes (Mongrain et al., 2010; Wisor et al., 2002).

To identify upstream regulatory elements that could explain these sex specific differences, transcription factor binding analysis using CHEA Transcription Factor Targets dataset was conducted. This analysis showed that binding sites for EZH2 (Enhancer of zeste homolog 2) and SUZ12 were significantly enriched in males (Figure 2C), and included transcripts such as *Adcy7, Bdnf* and *Fosl2* which are all upregulated only in males. EZH2 is a histone methyltransferase which binds to regulatory subunits, one of which is SUZ12, to form the polycomb repressor complex 2 (PRC2) which plays a role in gene silencing (Blackledge & Klose, 2021). EZH2 has multiple binding partners through which it binds to DNA, one of which includes E2F6. Inspection of the promoter regions using ENCODE Registry of candidate cis-Regulatory Elements (cCREs) and JASPAR Transcription Factor Binding Site Database from USCS browser identified E2F6 binding sites in both BDNF and Adra1B promoters (Table 1, Figure 2). Similar analysis revealed both AR (Androgen receptor) and ESR1/2 (Estrogen receptor 1/2) binding sites in both BDNF and Adra1B promoters (Table 1).

**Table 1:** Key transcription factor binding sites for BDNF and Adra1B. AR (Androgen receptor); ESR1/2 (Estrogen receptor 1/2).

| Gene | Transcription Factor binding | Binding site(s) |
| --- | --- | --- |
| BDNF | AR | chr2:109710346-109710362; chr2:109678105-109678121 |
|  | ESR1 | chr2:109694088-109694104 |
|  | ESR2 | chr2:109694089-109694103 |
|  | E2F6 | chr2:109695859-109695871 |
| *Adra1B* | AR | chr11:43826868-43826884; chr11:43776498-43776514 |
|  | ESR2 | chr11:43836892-43836906 |
|  | E2F6 | chr11:43823618-43823630; chr11:43837514-43837526 |

To understand the functional relevance of the genes that demonstrated sex-specific changes in their response to sleep deprivation, we conducted pathway analysis on the Enrichr platform (Chen et al., 2013), from which Bioplanet 2019 results are reported in Figure 2D. BDNF, HIF and NOS signalling are all significantly different between the sexes. STRING analysis to generate Protein-Protein Interaction Networks and to assess functional enrichment within connected networks (Figure 3). We found that the network had a hub at JUN which linked to a cluster containing BDNF, EGR3, FOSB, FOSL2, and HSP1A, all of which are upregulated in males but not females (See Figure 1A and B). Other clusters included sex-chromosome specific genes (FDR = 0.006, Figure S1A), vascular endothelin signalling genes (FDR = 0.01, Figure S1B), and collagen-containing extracellular matrix genes (FDR = 0.0001, Fig S1C) with FN1 (Fibronectin 1) at the hub, a gene which is upregulated only in males.

**Figure 3:**
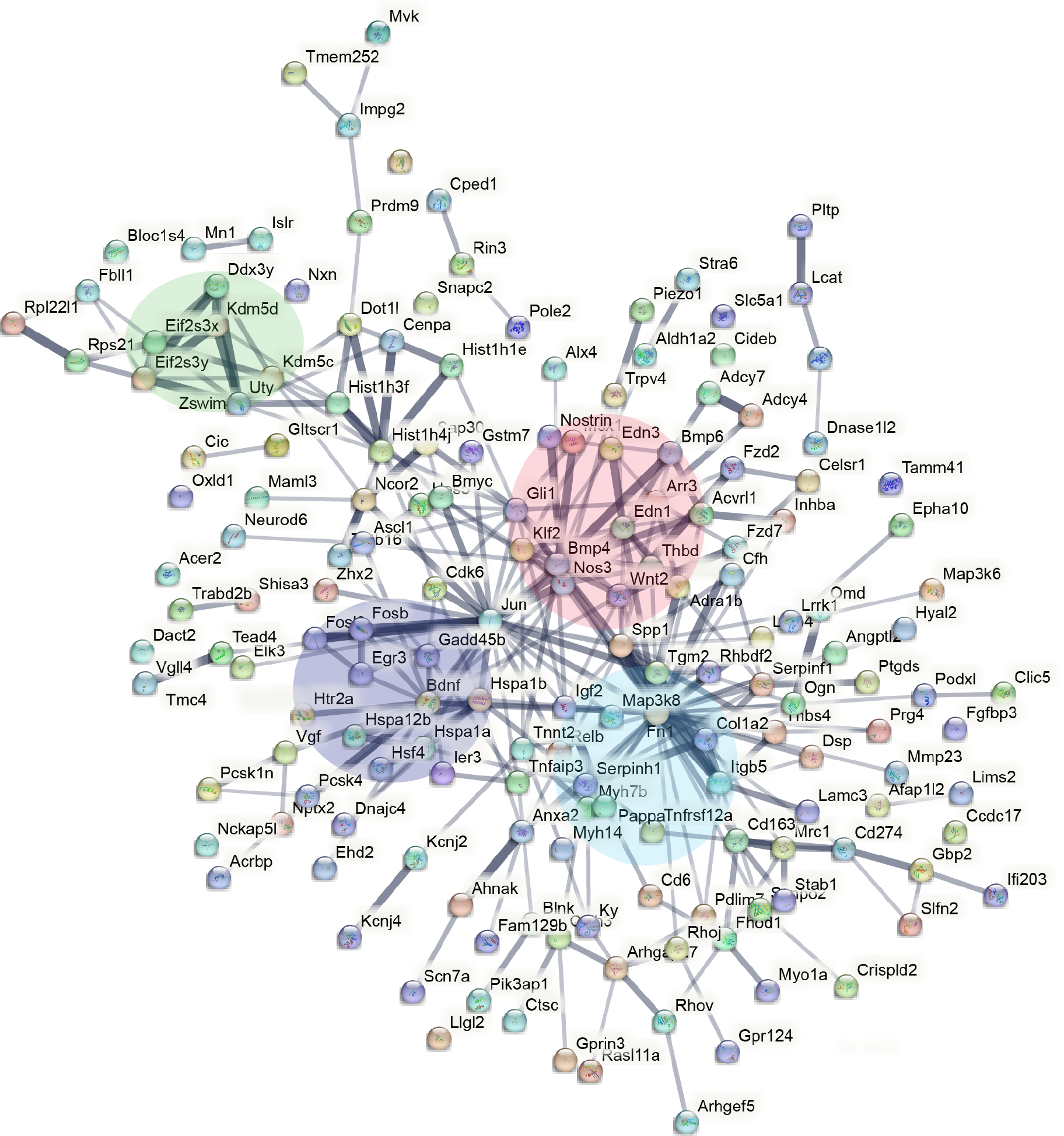
STRING interaction network mapping of the transcripts in Fig. 2A showing clusters centering around BDNF signalling (purple), sex-chromosome specific genes (green), endothelin signalling (red), and FN1/extracellular matrix (blue),.

## DISCUSSION

The molecular mechanisms that underlie sex differences in sleep phenotypes remain largely unknown. To this end, our studies show that there are subtle, but important differences in male and female cortical transcription profiles following acute sleep deprivation in mice. Most of the genes long-established to be differentially regulated following sleep deprivation, such as IEG *DBP*, demonstrate no sex difference (Figure 1A). However, a small number of genes show differential regulation in only one sex, such as *FOSL2*. These genes appear to cluster around *BDNF, JUN, FN1*, and *BMP4* in the protein-protein interaction network predicted by STRING.

The first of the molecular pathways differentially regulated in both sexes following sleep deprivation involves BDNF signalling. BDNF has neurotrophic plasticity roles (Bramham & Messaoudi, 2005) related to neuronal survival, learning, memory, and sleep (Bathina & Das, 2015; Giese et al., 2014). Unsurprisingly, reduced levels are associated with neurodegenerative diseases such as Alzheimer’s disease and Parkinson’s disease, but also depression and stress-related mental disorders for which sleep problems are a common symptom. This study shows that transcription of BDNF is increased significantly in males exclusively following sleep deprivation, suggesting the role of BDNF is reduced in females which would affecting synapse plasticity. This differential signalling mechanism offers a rationale for the observed sex-difference in which female rodents experience a greater impact on memory and cognitive performance due to sleep deprivation compared to males (Fernandes-Santos et al., 2012; Hajali et al., 2012).

Bmp4 (bone morphogenetic protein 4) serves as the hub for a cluster of genes related to endothelial signalling. Many elements of this pathway are differentially regulated in a sex-specific manner after sleep deprivation (Figure S1B). The gene *Acer2* (alkaline ceramidase 2) is upregulated in females only following sleep deprivation (Figure 1B and S2B), it encodes an enzyme that hydrolyses ceramides to generate the bioactive lipid sphindosine-1-phosphate (S1P) (F. Li et al., 2018). S1P regulates HIF-1a activity (Kalhori et al., 2013) and is known to be a major regulator of endothelial function (Figure S2A). These findings suggest that lipid signalling is different in males and females and given the growing evidence demonstrating a correlation between lipid profile and amount of sleep (Kaneita et al., 2008), the upregulation of Acer2 exclusively in females suggests an interesting sex-specific role of S1P signalling in regulating sleep.

A major hub of the network that is regulated by sleep in a sex-specific manner is Fibronectin 1 (*FN1*), which regulates extracellular matrix assembly and function (Parisi et al., 2020). This is surprising as not much is known about its role in sleep regulation, although some evidence points to the dysfunction of extracellular matrix molecules as part of the mechanism underlying synaptic dysfunction in psychiatric disorders and memory consolidation during sleep (Gisabella et al., 2021). Overall, further investigation is required to establish the functional relevance of these findings.

Similarly, not much is known about the role of ADRA1B in sleep regulation. Using linkage analysis and whole-exome sequencing, a mutation in ADRA1B gene has been identified in affected people with familial natural short sleep 2 (FNSS2) who have a lifelong reduction in sleep duration without suffering the cognitive consequences of sleep deprivation (Shi et al., 2019). Results of subsequent mice studies suggest a role of ADRA1B in activating neurons during REM sleep and wakefulness. Our study shows that expression of the normal gene is upregulated in females following sleep deprivation, it would be interesting to determine if the mutated form exhibits any sex-bias and whether this affects the phenotype of women with FNSS.

Through the promoter analysis, a network can start to be formed with EZH2 playing an important role at the top of the pathway. A study has shown that, separate to its PRC- and methylation-roles, EZH2 binds directly to the Ar (Androgen receptor) promoter and acts as a transcriptional activator of the Ar gene (J. Kim et al., 2018). Ar is a nuclear sex hormone receptor, which along with ESR1/2 (estrogen receptors 1/2), mediate downstream signalling events that control gene expression (Fuentes & Silveyra, 2019). JASPER promoter analysis identified multiple binding sites for these transcription factors in the promoter of BDNF and ADRA1B. Since EZH2 recruits Ar directly, enrichment of EZH2 binding sites following sleep deprivation correlates with male-only increase in expression of genes, such as BDNF, for which both Ar and EZH2 can act as a transcriptional activator. In support, we identified binding sites for Ar and EZH2 in key genes that were regulated in sex-specific manner (Table 2). Interestingly, a study investigating the sex differences in liver transcription programs showed that EZH2 also plays a key transcriptional role leading to sex-bias in susceptibility to fibrosis and other liver diseases (Lau-Corona et al., 2020). While this supports our identification of EZH2 as a potential mediator of sex-specific transcriptional programs in response to sleep deprivation, the exact influence of this transcription factor requires further investigation.

This study is limited by the fact that we have not carried out any functional validation of the transcriptomic changes. Furthermore, transcriptomic changes are not necessarily reflected at the level of the protein, where translational and post-translational regulation can significantly alter the outcomes from transcriptional changes. Nevertheless, the goal of this study was to provide a description of transcriptional pathways that would inform future studies. It revealed some intriguing molecular pathways that are differentially regulated in both sexes following sleep deprivation that may provide important insight into the mechanisms that underlie sleep regulation and the response to loss of sleep. Additionally, this data describes, for the first time, how male and female mice react differently at the transcriptomic level to sleep deprivation. These small subtle differences could have big functional implications and shows that females process the signals from sleep deprivation differently, with the identified genes providing a groundwork for further investigation to uncover the molecular pathways that underlie sex differences in sleep regulation. To conclude, although traditionally neuroscience research is conducted exclusively in males, this study has shown that many of the key transcripts associated with sleep deprivation respond similarly in both sexes with differences not widespread and therefore can be easily factored into analysis. This has strengthened the case for the inclusion of females in future research.

## FIGURE LEGENDS

**Figure S1:**
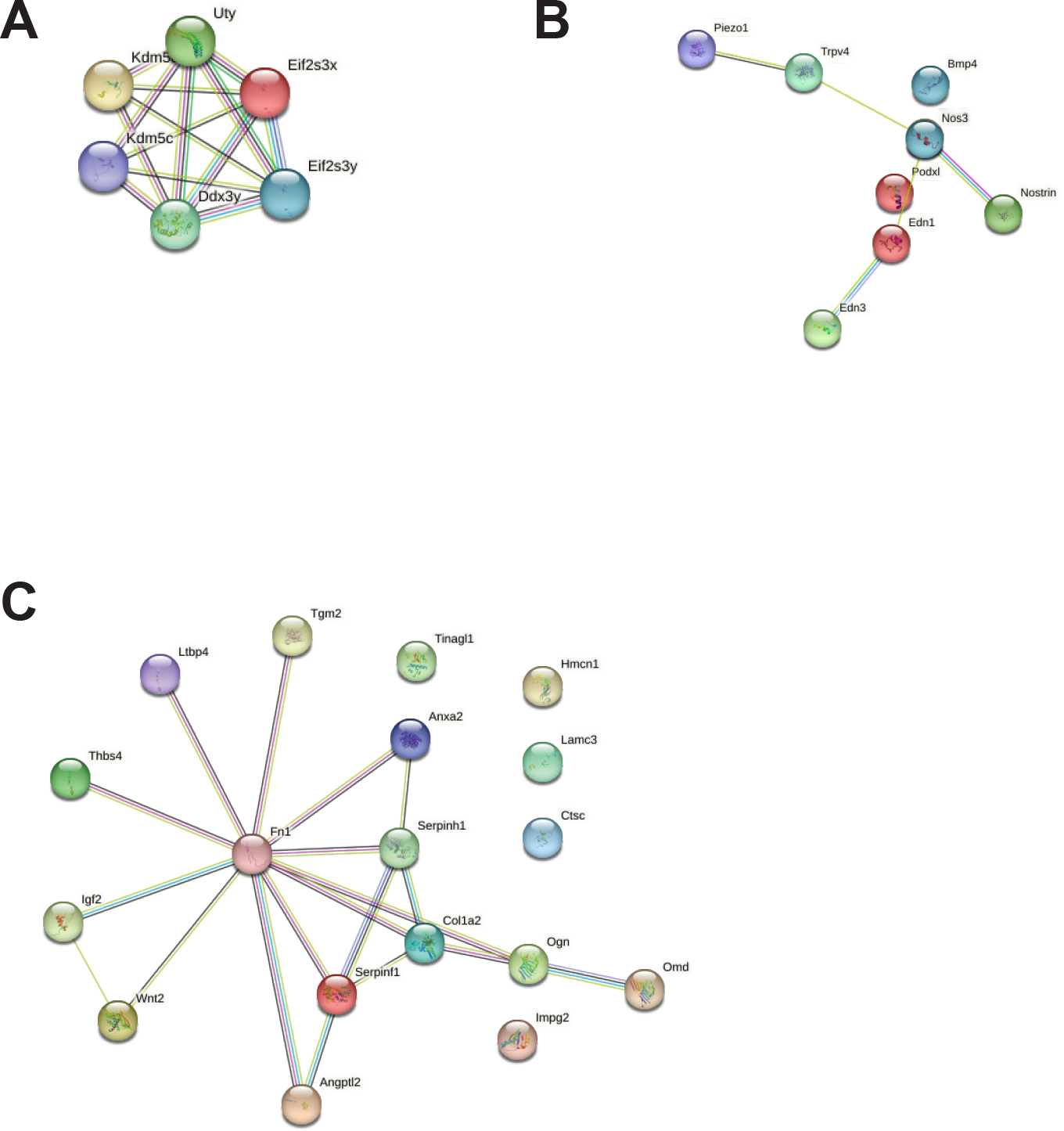
Related to Figure 3. Specific functional networks from STRING analysis for (A) sex-chromosome specific genes, (B) endothelin signalling and (C) extracellular matrix.

**Figure S2:**
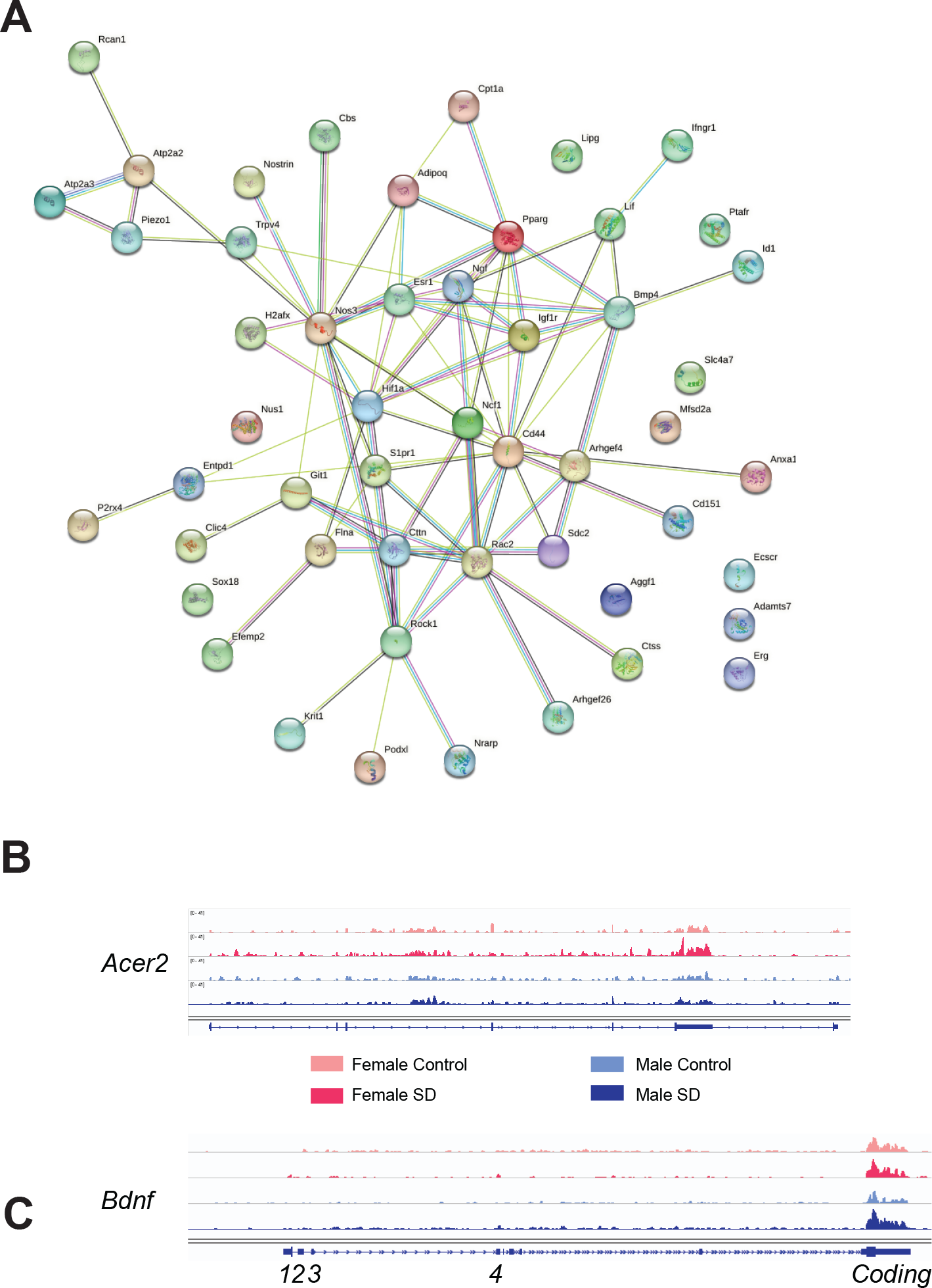
Related to Figure 1-3. (A) The entire endothelin signalling network. RNA-Seq tracks under different conditions as labelled for (B) *Acer2* and (C) *Bdnf*, with specific exons labelled.

## ACKNOWLEDGEMENTS

This work was supported by the following sources of funding: BB/N01992X/1 David Phillips fellowship from the BBSRC to AJ, and Oxford-Elysium Cellular Health Fellowship to LT.

## AUTHOR CONTRIBUTIONS

LT and QD conducted the experiments. TS, LT, QD and IS and AJ analysed data. TS, IS and AJ co-wrote and edited the manuscript, with input from all authors.

## DECLARATION OF INTERESTS

AJ is a founder Circadian Therapeutics Ltd. and hold shares in the company.

